# *De novo* meristem development in *Marchantia* requires light and an apical auxin minimum

**DOI:** 10.1101/2025.07.17.665278

**Authors:** Eva-Sophie Wallner, Natalie Edelbacher, Liam Dolan

## Abstract

Meristems are generative centres with stem cells from which the bodies of land plants develop. *Marchantia polymorpha* spores are single cell structures formed at meiosis. On germination, spores divide asymmetrically to form a basal cell that terminally differentiates and an apical germ cell that divides into an early cell mass on which a flat prothallus develops. A single stem cell niche (meristem) forms *de novo* at the margin of the prothallus to drive development of the thallus plant body. Here we show that the prothallus forms at the apical pole of the early cell mass and represses the formation of other prothalli. LOW AUXIN RESPONSIVE (MpLAXR) marks this apical pole indicating that an auxin minimum is located at the site of organogenesis. Light is required for the formation of the apical auxin minimum and for the development of the prothallus from the early cell mass. Disrupting the apical auxin minimum by exogenous auxin treatment suppresses the transitions to the prothallus and formation of the meristem from the early cell mass. A similar molecular program operates during plant regeneration from a single differentiated thallus cell, which regains stemness (pluripotency) upon surgical isolation from surrounding tissues; the isolated cell divides forming an early cell mass that develops a local auxin minimum where a flat prothallus with a single meristem forms. We conclude that a light-dependent, apical auxin minimum is required for the formation of the prothallus and the *de novo* development of the first meristem in *Marchantia polymorpha*.

**Graphical abstract:** 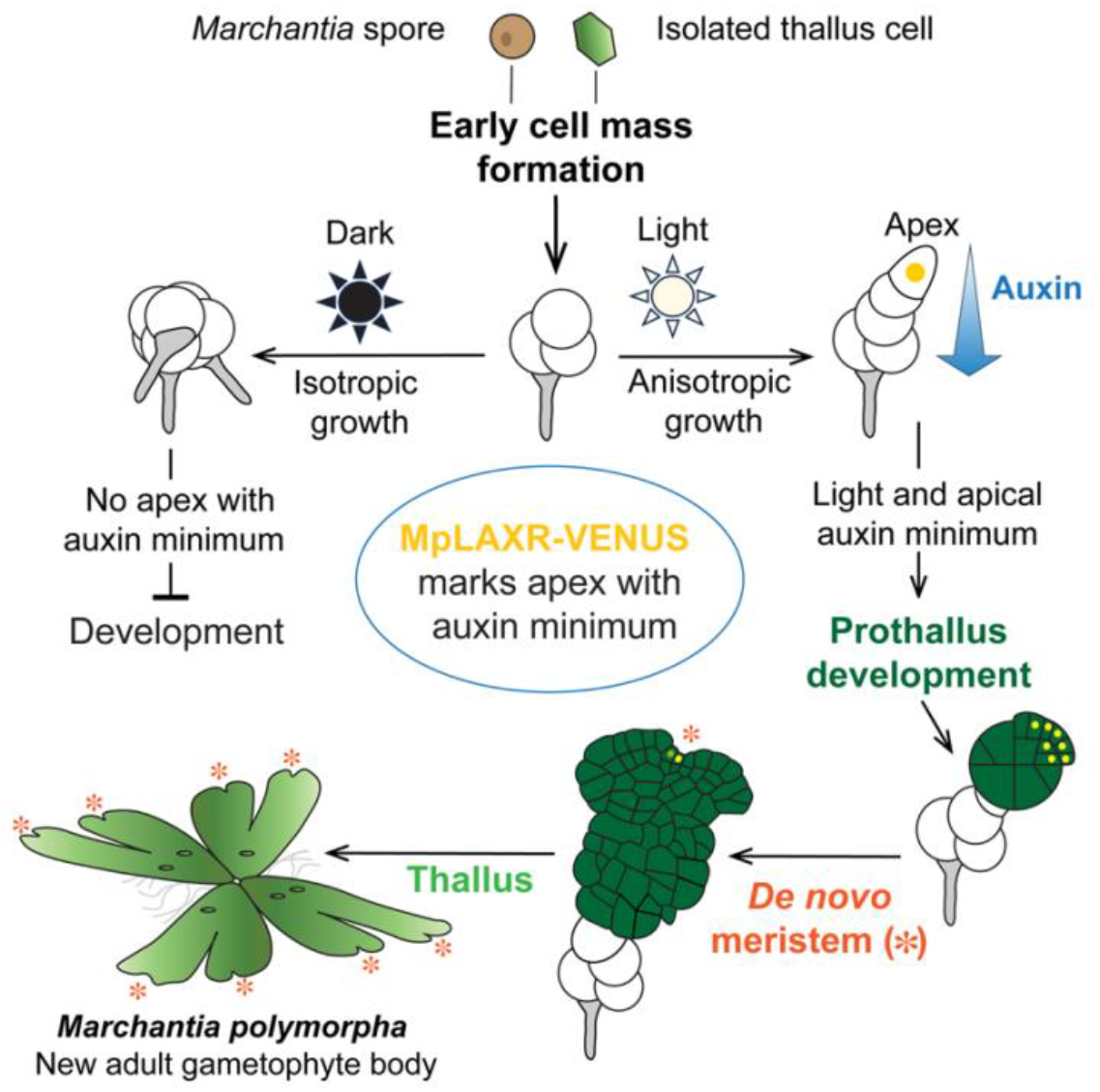

## INTRODUCTION

Plant meristems contain stem cell niches from which the multicellular body develops. The multicellular gametophyte plant body of the liverwort *Marchantia polymorpha* (*Marchantia*) develops from haploid spores^1,2^. Spores are single cells and products of meiosis that develop in isolation from the parental plant^2^. The first division of the germinated spore is asymmetric and forms a relatively large apical germ cell with proliferative potential and a relatively small basal cell that differentiates as a germ rhizoid cell^3-7^ (Figure 1A). The position of the germ rhizoid cell defines the location of the basal pole and the apical germ cell the location of the apical pole (Figure 1A). The apical germ cell divides forming a population of largely isotropically growing cells referred to as the early cell mass, or as the protonema^1,4,8^ (Figure 1A). The morphology of the early cell mass can vary but it eventually produces a prothallus – a flat structure with dorsoventral polarity.

**Figure 1:**
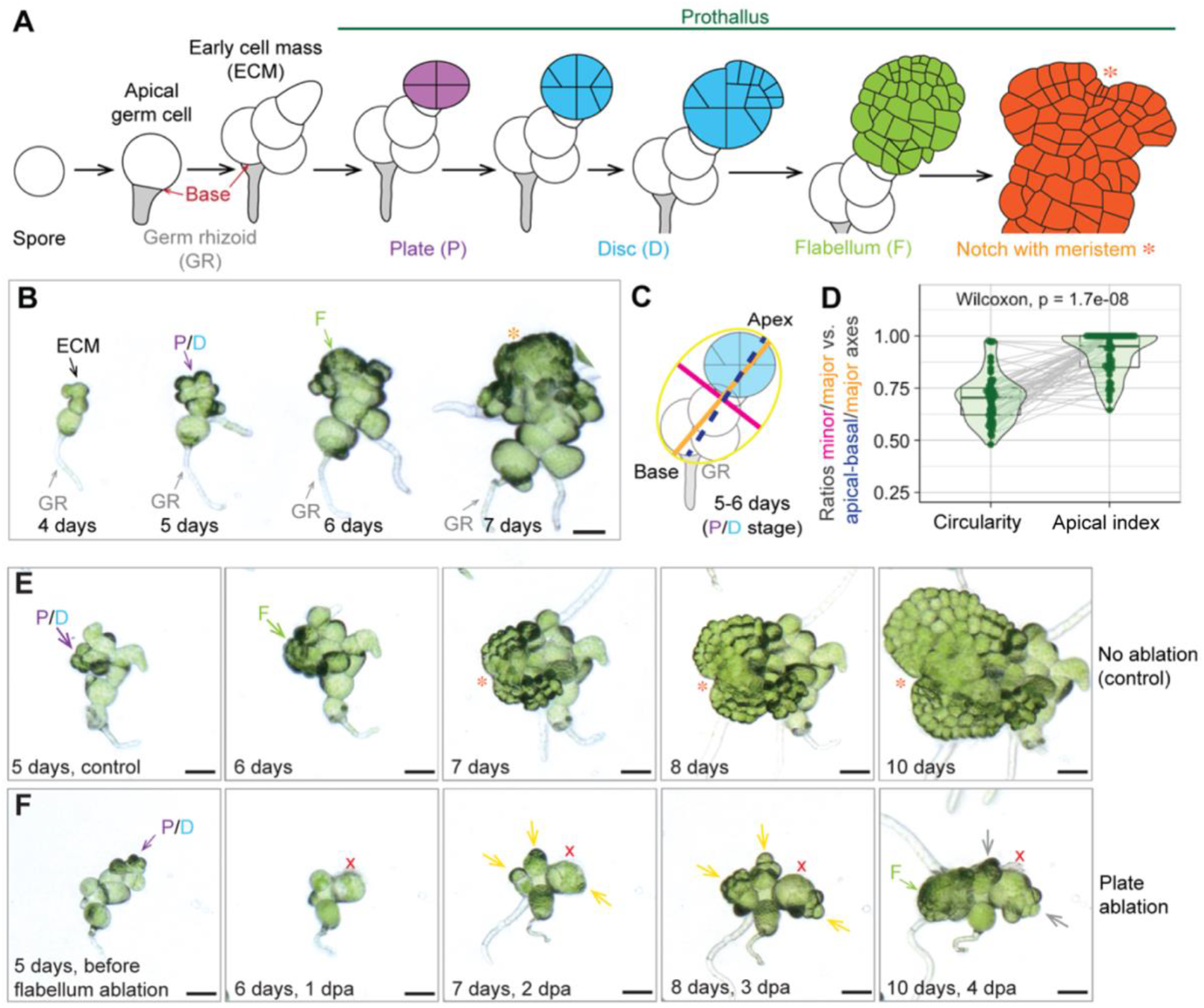
The prothallus forms at the sporeling apex and represses formation of additional prothalli. **A**) Schematic representation of *Marchantia polymorpha* (*Marchantia*) development from a single-celled spore. After proliferation of the apical germ cell into an early cell mass, the prothallus forms from an early cell mass cell via reproducibly oriented divisions^8^ to give rise to the first *de novo* formed meristem (marked by asterisks). **B**) Time course experiment of Tak-1xTak-2 (wild type) sporelings from day 4-7 as seen in bright field microscopy and when grown in standard continuous light conditions (45 µmol m^-1^ s^-2^). The prothallus formed at the sporeling apex and was positioned distally to the germ rhizoid (GR). The GR indicates the start point of development (base). The distance between sporeling apex and GR defines the apical-basal body axis. Scale bar 50 µm. **C**) Prothallus positioning was quantified in 5-to 6-day-old sporelings that had formed a young prothallus in the plate or disc stage. An ellipse was fitted to encompass all green sporeling cells. The lengths of the major (orange line) and the minor axis (pink line) as well as the apical-basal body axis (blue dashed line) were determined. **D**) Statistical analysis of prothallus positioning with respect to overall sporeling shape as described in C. For each sporeling (n = 50, repeated twice) the measure of circularity (minor over major axis) was compared to its apical index (apical-basal over major axis) by a paired Wilcoxon test with α = 0.05. The apical index was significantly greater than the measure of circularity (p-value = 1.7e-08), indicating that the apical-basal axis aligned closely with the major axis of the sporeling and hence sporeling growth is anisotropic with a defined start point (base) and end point (apex). **E-F**) Ablation of a young prothallus in the plate or disc stage (red x) led to cell proliferation at other sites (yellow arrows), of which one developed into a flabellum (green arrow in E), while proliferation at the other sites stopped (grey arrows). n = 93 in 2 independent experiments. Unablated sporelings served as control (D). Scale bars 50 µm.

Previously, we defined five developmental stages based on characteristic cell division patterns during prothallus development^8^. The first morphologically distinct phase of prothallus development is the plate stage. A single cell on the surface of the early cell mass that we designated as prothalloblast undergoes first a symmetric cell division followed by two divisions that are oriented perpendicular to the first division, thus forming a quadrant of cells^4,8^ (Figure 1A). Oblique divisions in three of the four plate quadrants produce daughter cells around the margin that expand the structure into a multicellular disc^8^. Cells derived from one of the quadrants divide more than the others and contribute disproportionately to the formation of a flat fan-like structure we designated as a flabellum^8^. Cell divisions in the flabellum become gradually restricted to a small marginal region to form a sinus flanked by two lobes^1,8,9^. The first meristem develops at the base of the sinus and together these constitute the meristematic notch^1,9^. An apical stem cell in the meristem divides on four cutting faces to form dividing populations of cells that develop thallus axes comprising tissue in three dimensions^2,9^. Meristems duplicate periodically forming a series of bifurcating axes^1,2,8,9^. However, we know neither how the site of prothallus initiation is positioned on the early cell mass nor how the site of the first meristem is positioned on the prothallus (Figure 1A).

*Marchantia* also has the capacity to regenerate meristems from differentiated cells without the addition of hormones or dedifferentiation factors. For example, meristems develop from differentiated cells in isolated thallus fragments after surgical removal of the active meristem^10,11^. Furthermore, individual differentiated cells divide and regenerate meristems when isolated from neighbouring cells by surgery or after protoplasting^12-14^. However, we do not know how internal signals guide meristem generation in sporelings or meristem regeneration when differentiated cells are reprogrammed.

The ethylene response factor *LOW AUXIN RESPONSIVE* (Mp*LAXR*) – also known as Mp*ERF20* (Mp5g06970) – specifically marks the apical stem cell within the Marchantia meristem^15^. Mp*LAXR* promotes stemness and – if ectopically expressed – induces dedifferentiation of tissues, followed by cell division and meristem formation^10,15^. In dissected thallus pieces, Mp*LAXR* is expressed at the cut-site and promotes cell division and meristem regeneration^10^. The Mp*LAXR* promoter is activated if auxin levels are low and is repressed by auxin treatments^10^. Auxin is a plant hormone and predicted morphogen that controls diverse aspects of plant development^16^. It is hypothesised that auxin levels are low in apical cells of *Marchantia* meristems and that this auxin minimum is important for maintenance of stem cell function^10,17^. These localised auxin minima are predicted to result, at least in part, from the activity of PIN auxin efflux carriers in surrounding cells that transport auxin away from the apex^17^. Experimental treatment of regenerating meristems with exogenous auxin both blocks meristem regeneration from excised thallus axes and eliminates wounding-induced Mp*LAXR* expression at the cut-sites of thalli^10,14^. We therefore propose that Mp*LAXR* can be used as a marker to image auxin minima during meristem development in *Marchantia*^14,15^.

Here we show that the Marchantia sporeling develops an apical-basal axis with an auxin minimum at the apical pole. This apical auxin minimum forms at the apical region of the early cell mass and is propagated in the developing prothallus where it becomes progressively restricted to the single apical stem cell of the meristem. Light is required for developmental progression and establishment of the apical auxin minimum. Disruption of the auxin minimum also blocks prothallus development and meristem initiation. We also show that plants regenerating from single, isolated differentiated thallus cells, also develop an apical auxin minimum associated with the initiation of prothallus organogenesis. These data suggest that light initiates the formation of a local auxin minimum required for *de novo Marchantia* meristem development in sporelings. We propose that a similar mechanism operates during meristem generation in the sporeling and meristem regeneration from differentiated cells in the thallus.

## RESULTS

### The prothallus forms at the sporeling apex and represses formation of additional prothalli

The *Marchantia* prothallus develops from a single cell – the prothalloblast^8^ – that forms 3-4 days after spore germination on the surface of the early cell mass (Figure 1A)^8^. The shape and cell number of the early cell mass varies^1,4,8^. We therefore asked if the overall shape of the early cell mass could predict the site of prothallus initiation or if prothallus positioning on the early cell mass was random. To distinguish between these alternatives, we performed time-course imaging on 50 randomly chosen sporelings from the early cell mass stage (day 4) to the meristem stage (day 7 marked with an orange asterisk) (Figure 1B). Formation of the four-celled plate or seven-celled disc indicated that the prothallus had initiated – typically by day 5 (Figure 1B). To quantitatively define the shape of the early cell mass, we approximated these 5-day-old sporelings with an ellipse that encompassed all cells except the rhizoid (see yellow outline that excludes rhizoids in Figure 1C). Each ellipse is defined by two axes of symmetry: the major axis represents its longest diameter (marked in orange), and the minor axis represents its shortest diameter (marked in magenta) (Figure 1C). The ratio of the minor axis to the major axis is a measure of circularity or anisotropy of the sporeling. A ratio of 1 describes a perfect circle, while a ratio closer to 0 describes an anisotropic (elliptical) sporeling shape (Figure 1C-D). Our data revealed that all sporelings were elliptical and none fitted into a perfect circle, suggesting that the early cell mass grew anisotropically along the major axis, from sporeling base towards apical pole (Figure 1C-D).

We next asked if the prothallus originated near the apical pole of the anisotropically growing early cell mass or emerged at random positions on the surface of the early cell mass. To distinguish between these alternatives, we determined the distance between the sporeling base and apex (Figure 1C). The sporeling base is defined by the point where the germ rhizoid attaches to the early cell mass (Figure 1A). The prothallus initiation site marks the future plant apex, where the meristem is eventually formed (orange asterisk in Figure 1A). The distance between base and apex therefore defines the apical-basal axis of each sporeling (dashed blue line). To determine how close the prothallus initiation site was to the apical pole of the early cell mass, we calculated the apical index, which is expressed by the ratio of the length of the apical-basal axis (dashed blue line) to the length of the major axis (orange line) of the ellipse that circumscribed the sporeling (Figure 1C). A ratio of 1 represented the greatest possible distance between apex and base, meaning that germ rhizoid and prothallus were positioned at opposite ends of the early cell mass. A ratio closer to 0 indicated that prothallus initiation and germ rhizoid base were close to each other (Figure 1C-D). The observed ratios ranged from 0.6–1, indicating that the prothallus initiation site was always positioned closer to the apical pole of the early cell mass than to the sporeling base (Figure 1B-C). Finally, we compared each sporeling’s circularity with its apical index. The apical index was significantly greater than the circularity (Figure 1D) and sporelings with the lowest circularity had the highest apical index, as seen by the trend of grey lines in the paired analysis plot (Figure 1D). This suggested that the more anisotropically the early cell mass grew, the closer the prothallus initiated at its apical pole. We conclude that anisotropic early cell mass growth promotes prothallus positioning at the apical pole, consequently leading to prothallus development along the apical-basal growth axis of the sporeling.

We typically observed a single site of prothallus initiation from the early cell mass (Figure 1B-D). Therefore, we hypothesized that development of the prothallus at the early cell mass apical pole repressed initiation of prothalli elsewhere on the early cell mass. To test this hypothesis, we ablated the young prothallus at the plate or disc stage in 5-day-old sporelings with a 355 nm DPSS laser and tracked sporeling development after the ablation (Figure 1E-F). Ablating the young prothallus initiated new cell division in several positions on the early cell mass within two days of ablation (dpa) (yellow arrows in Figure 1F). Within three days, new cell clusters emerged at the sites of these divisions (yellow arrows in Figure 1F). Four days after ablation, a new prothallus developed from one of these cell clusters while no other clusters of the early cell mass developed further (Figure 1F). These data are consistent with the hypothesis that the first prothallus represses the development of other prothalli in the sporeling.

### The first meristem develops at the central flabellum margin

The first *Marchantia* meristem is located in a notch – a region that includes a sinus between lobes of expanded prothallus – that emerges in sporelings between day 7 and 10^2,8^. We previously designated this expanded prothallus stage as flabellum because of its fan-like shape^8^. To define the position of the notch within the flabellum, we determined the flabellum midpoint between day 7 and 8, when the notch was first detectable (Figure 2A-B, green line in diagram at basal flabellum side) and calculated the acute angle from that midpoint to the notch (Figure 2B, orange line in diagram). Most data points ranged from 50° to 125° as depicted in the violin plot and the average notch angle did not significantly differ from 90° (Figure 2C). This indicated that the notch formed most frequently at the apex of the convexly curved flabellum margin (Figure 2A-B). The flabellum apex is the region along the flabellum margin that is perpendicular to the flabellum base (Figure 2B). We therefore reasoned that the number of notches may be correlated with the number of flabella that formed during sporeling development. We sampled developmental time courses of 100 sporelings. 97 sporelings developed a single flabellum that formed one meristem (Figure 2A and Figure 2E). Two sporelings developed a pair of adjacent flabella (Figure 2D-E). Whenever two flabella developed, the notch with a single meristem formed on the larger of the two flabella and was asymmetrically displaced to the side, nearer the second flabellum (Figure 2E). This indicates that a single meristem forms in the sporeling, even when two flabella develop. We never observed more than two flabella at this developmental stage of the sporeling. The dominance of one flabellum over others suggests that mobile signals could control the flabellum number and the positioning of meristems in sporelings.

**Figure 2:**
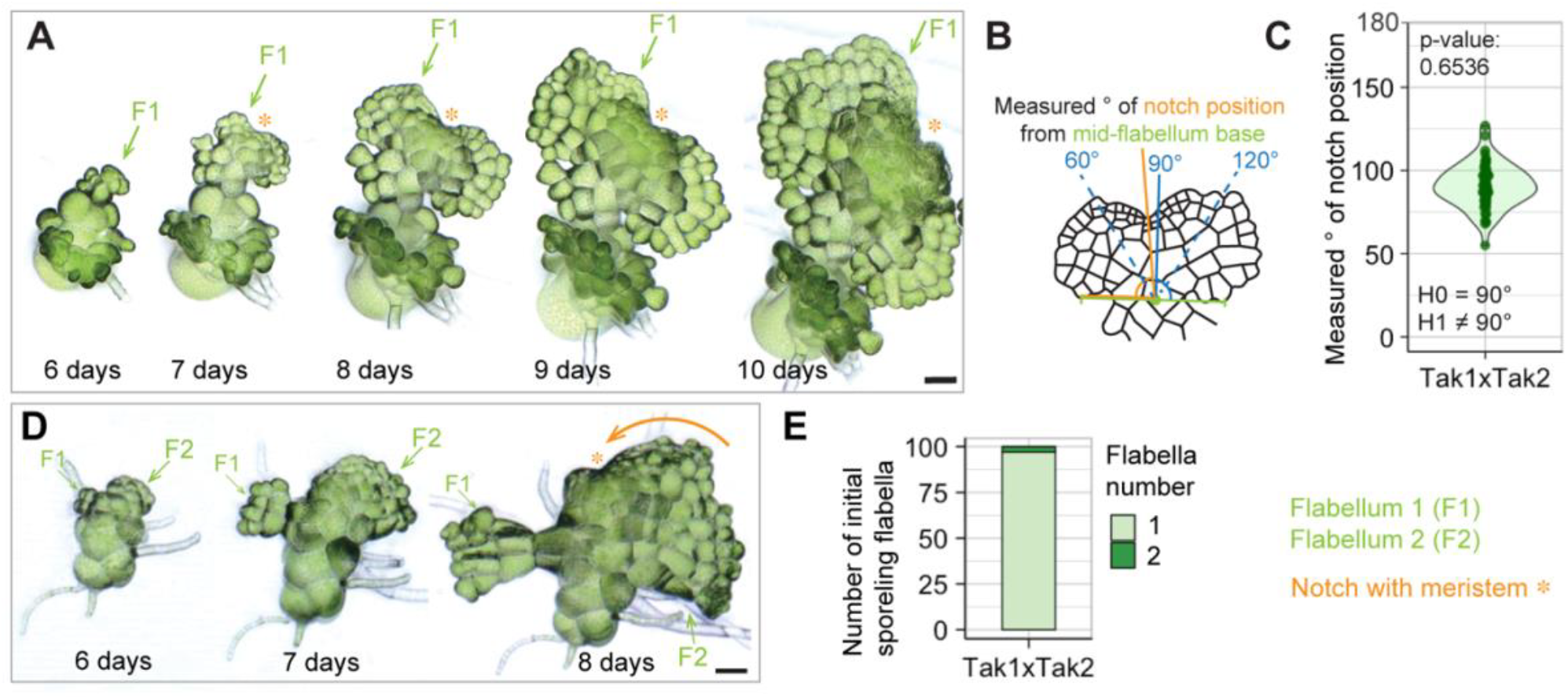
The first meristem develops at the central flabellum margin. **A**) Time course imaging of Tak-1xTak-2 (wild type) sporeling development between days 6-10. A flat flabellum (light green arrow) developed around day 5 and expanded throughout day 10. The notch, characteristic for *Marchantia* meristems, developed around day 7 (orange asterisk). Scale bar 50 µm. **B-C**) Statistical analysis of notch positioning in 8 day-old sporelings grown in standard CL with n = 50 (2 repetitions). The midpoint of the flabellum was determined (green line in B) and the acute angle to the notch with meristem calculated (orange line in B). A one-sample t-test for notches positioned at 90° on average (H0) or at angles other than 90° (H1) revealed with a p-value of 0.6536 and a mean of 90.86 that notch angles are on average positioned at a 90° angle. No notches were observed with angles greater than 130° or smaller than 50°. **D**) Time course imaging of a Tak-1xTak-2 (wild type) sporeling with two flabella (F1 and F2), with an off-centre positioning of the notch (orange asterisk) on the bigger flabellum (F2) but skewed towards F1. Scale bar 50 µm. **E**) Quantification of sporeling flabella within n = 100 individual sporelings. 98% produced only one flabellum, while 2% developed two flabella. Three independent repetitions with n = 100 each showed only one flabellum in 100% of the population, indicating that formation of two flabella is a rare event in Tak-1xTak-2 (wild type) sporelings.

### MpLAXR marks the sporeling apex throughout all stages of prothallus development

We hypothesised that an apical auxin minimum formed during both flabellum development and meristem formation and that these auxin minima would be marked by Mp*LAXR* (Mp5g06970)^10,14,15^. If our hypothesis were correct, we would expect MpLAXR protein to accumulate in restricted regions – auxin minima – along the flabellum margin before meristem formation. To test this, we generated a transgenic *Marchantia* line expressing the *pro*Mp*LAXR:*Mp*LAXR-VENUS* translational fusion (yellow signal) and *pro*Mp*UBE2:mCherry-g*Mp*RCI2*, which expressed the plasma membrane marker g*Mp*RCI2 (Mp3g18270.1) fused to the fluorescent protein mCherry (magenta signals). We imaged signal intensities of the respective fluorescent protein fusions in developing sporelings over the course of 10 days (Figure 3). At 24 hours, most germinated spores expressed the plasma membrane marker, but no MpLAXR-VENUS signal was detected (Figure 3A). No nuclear MpLAXR-VENUS signal was observed at the two-cell stage (48 hours) (Figure 3B). At day 3, MpLAXR-VENUS was present in the nucleus of a single cell at the apical pole of the early cell mass (Figure 3C). This cell gave rise to the prothallus plate (Supplementary video 1 and Figure 3D) in which MpLAXR-VENUS fluorescence was observed in all cells. MpLAXR-VENUS became restricted to the growing plate quadrant as the disc formed (Figure 3 E). MpLAXR-VENUS signal was present in most cells of the young flabellum before it became progressively restricted to the flabellum apex (Supplementary video 2 and Figure 3F-G). These data indicate that auxin minima develop transiently at developmental transitions that occur during prothallus development.

**Figure 3:**
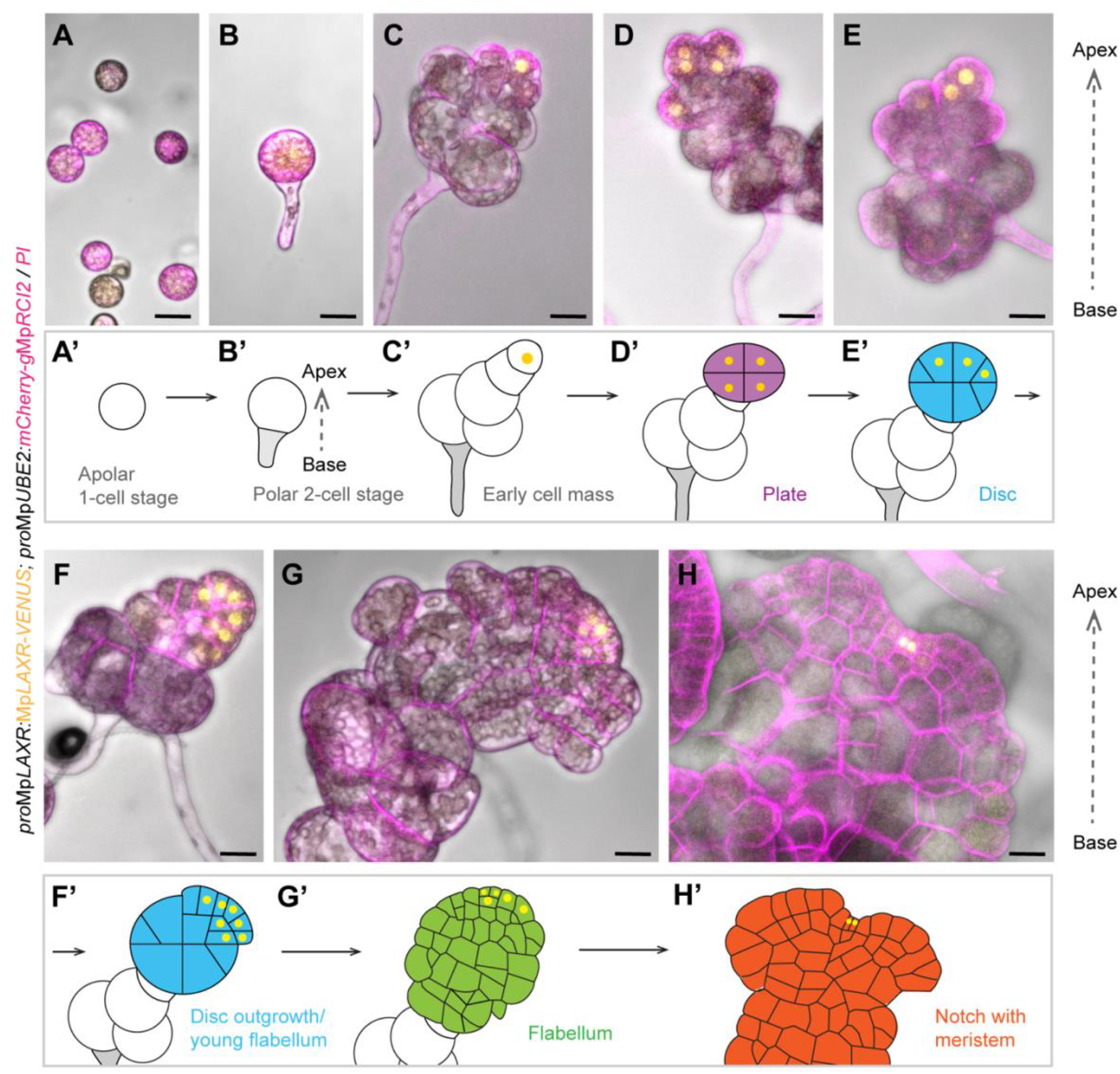
MpLAXR marks the sporeling apex throughout all stages of prothallus development. Spores expressing *pro*Mp*LAXR:*Mp*LAXR-VENUS* and *pro*Mp*UBE2:mCherry-*Mp*RCI2* were germinated and imaged at distinct developmental stages. Scale bars 20 µm. **A**) No MpLAXR-VENUS signal was observed in apolar single-celled spores 24 h post-plating. n = 163. **B**) At the 2-cell stage the apical-basal axis is defined by the apical germ cell with weak cytoplasmic and/or nuclear MpLAXR-VENUS signal and a rhizoid without MpLAXR-VENUS signal. n = 67. **C**) In the early cell mass and the prothalloblast, which signifies the first stage of prothallus development, MpLAXR-VENUS signal is typically restricted to the most apically positioned cell. n = 83. **D**) At the 2^nd^ prothallus stage, MpLAXR-VENUS signal can be detected in the nuclei of all four plate cells and occasionally in cells adjacent to the plate. n = 14 **E-F**) In the disc, MpLAXR-VENUS signals become restricted to the quadrant that proliferates and expands into the flabellum. n = 10. **G**) In the expanded flabellum, MpLAXR-VENUS signals are restricted to few cells at the flabellum apex. n = 63. **H**) After notch formation, MpLAXR-VENUS signals are only observed in 1-2 small and dividing apical cells of the flabellum margin that are embedded within the base of the notch. n = 38. **A’-H’**) Schematic representations of all major stages of sporeling and prothallus development with yellow signal summarising the observed MpLAXR-VENUS signals per stage and throughout 9 independent repetitions that yielded the sample sizes (n) per stage as listed above.

After the apical notch formed, the MpLAXR-VENUS signal was restricted to one or sometimes two apical stem cells of the meristem (Figure 3H). The single MpLAXR-VENUS-expressing cell divided to form two daughter cells, where the VENUS signal remained in one but gradually disappeared from the other daughter (Supplemantary video 2). We interpret the one cell that continuously expresses MpLAXR-VENUS as the apical stem cell of the meristem^9^. We conclude that a local auxin minimum, marked by MpLAXR-VENUS, is first established at the apical pole of the early cell mass, expands throughout the young flabellum before becoming restricted to the future apical stem cell at the flabellum apex. We therefore propose that the formation of a local auxin minimum precedes *de novo* meristem formation in *Marchantia* sporelings.

### Exogenous auxin treatment represses prothallus development and apical MpLAXR expression

We hypothesised that a local auxin minimum at the apical pole of the early cell mass was required for prothallus and meristem development. To test this hypothesis, we experimentally eliminated the apical auxin minimum by treating sporelings with 5 µM 1-naphtaleneacetic acid (NAA), a synthetic auxin that is not degraded by the plant’s endogenous peroxidase system^18^. As controls, plants were treated with the solvent DMSO without NAA (mock treatment). Sporelings grown on mock plates initiated small prothalli by day 4 (green arrow in Figure 4A), prothalli with a small flabellum by day 6 (green arrow in Figure 4B) and a notch with a single, active meristem at the flabellum apex by day 8 (green arrow in Figure 4C). Formation of the meristem by day 8 therefore marked the endpoint of prothallus development^8^. By contrast, sporelings germinated and grown on 5 µM NAA formed an early cell mass with multiple rhizoids but developed neither prothalli (plate, disc or flabellum) nor meristems by day 8 (Figure 4D-F). These data suggest that NAA-treatment suppressed the initiation of the prothallus and are consistent with the hypothesis that an apical auxin minimum is required for prothallus development.

**Figure 4:**
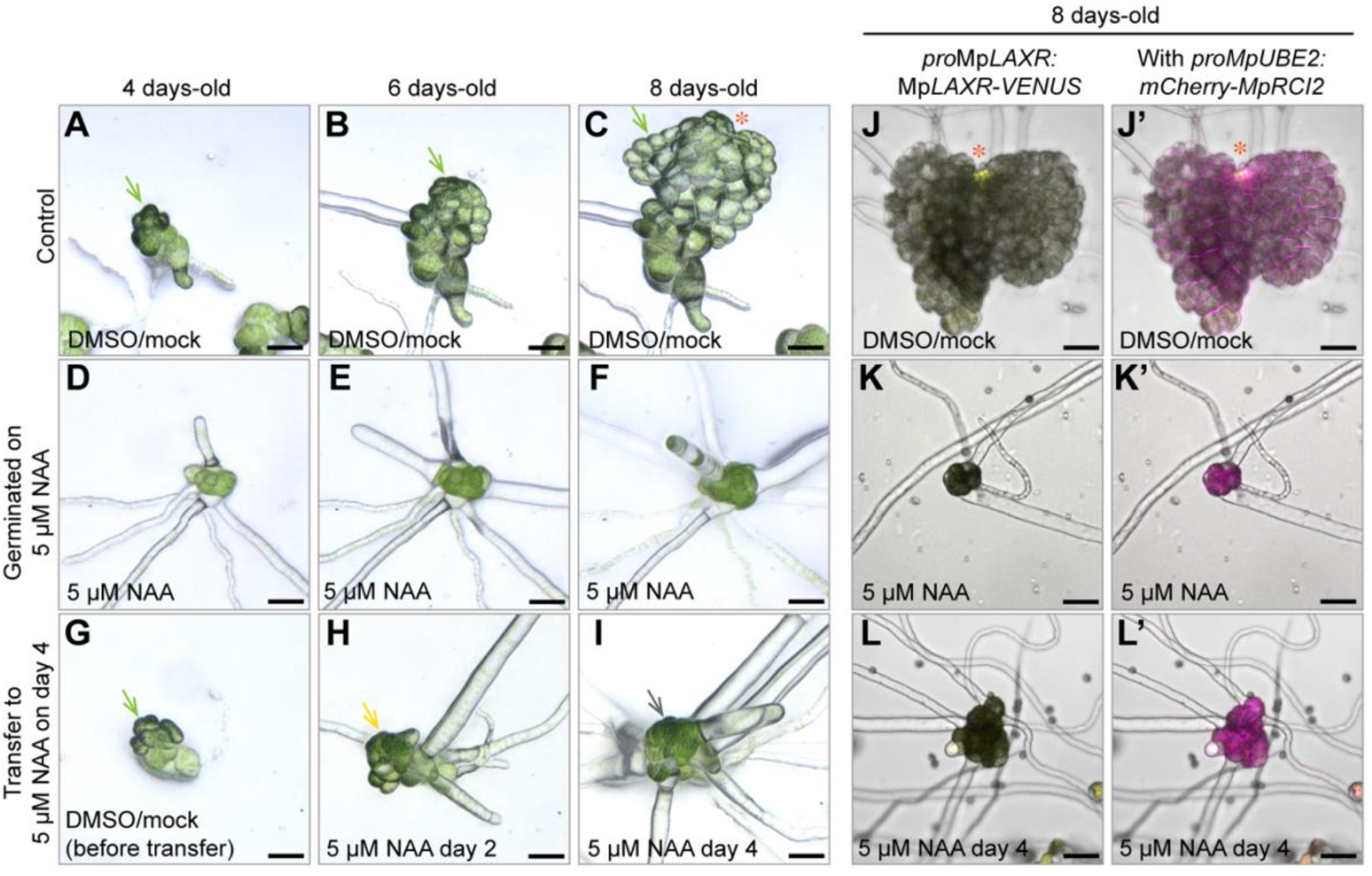
Exogenous auxin treatment represses prothallus development and apical MpLAXR expression. **A-C**) Tak-1xTak-2 (wild type) sporelings were grown on cellophane-covered standard media supplied with the solvent DMSO (mock) as control and their developing flat flabellum was imaged at day 4 (A, green arrow), day 6 (B, green arrow) and day 8 (C, green arrow). Sporelings tracked: n = 167 in 3 repetitions. **D-F**) Tak-1xTak-2 (wild type) sporelings were grown on cellophane-covered standard media supplied with 5 µM NAA and imaged at day 4 (D), day 6 (E) and day 8 (F). Sporelings tracked: n = 134 in 3 repetitions. **G-I)** Cellophane pieces with 4 day-old sporelings with tiny flabella (G, green arrow) grown on control media were transferred to media supplemented with 5 µM NAA, which slowed down (E, yellow arrow) and later aborted flabellum growth (F, grey arrow). Sporelings tracked: n = 81 in 2 repetitions. Scale bars 50 µm. **J-L**) Tak-1xTak-2 sporelings expressing *pro*Mp*LAXR:*Mp*LAXR-VENUS; pro*Mp*UBE2:mCherry-*Mp*RCI2* were grown for 6 days on standard media supplemented with the solvent DMSO/mock (J), transferred afer 4 days to media supplemented with 5 µM NAA (K) or germinated and grown for 6 days on media with 5 µM NAA (L). Split channels depict the MpLAXR-VENUS signal (J-L, yellow), which was only observed at the notch in the mock-treated control (J) and merged images with plasma membrane signal (J’-L’). n = 18 (J), 40 (K), 25 (L) in 3 repetitions. Scale bars 50 µm.

We next asked if auxin treatments impacted only prothallus initiation or also later prothallus development (Figure 1A)^8^. To answer this, we transferred 4-day-old sporelings, which had each developed a single small prothallus (green arrow in Figure 4G), from mock plates to 5 µM NAA-containing plates and tracked their development (Figure 4H-I). After 2 days on 5 µM NAA, cells started to grow isotropically (yellow arrow in Figure 4H). The isotropic cell growth increased with NAA exposure time. Furthermore, the flat prothallus cells that had developed during the first 4 days on plates without NAA, enlarged and became spherical and were morphologically indistinguishable from early cell mass cells after 4 days of NAA-treatment (grey arrow in Figure 5I). We conclude that externally applied NAA was sufficient to suppress development of the prothallus and may revert prothallus cells to early cell mass cells.

**Figure 5:**
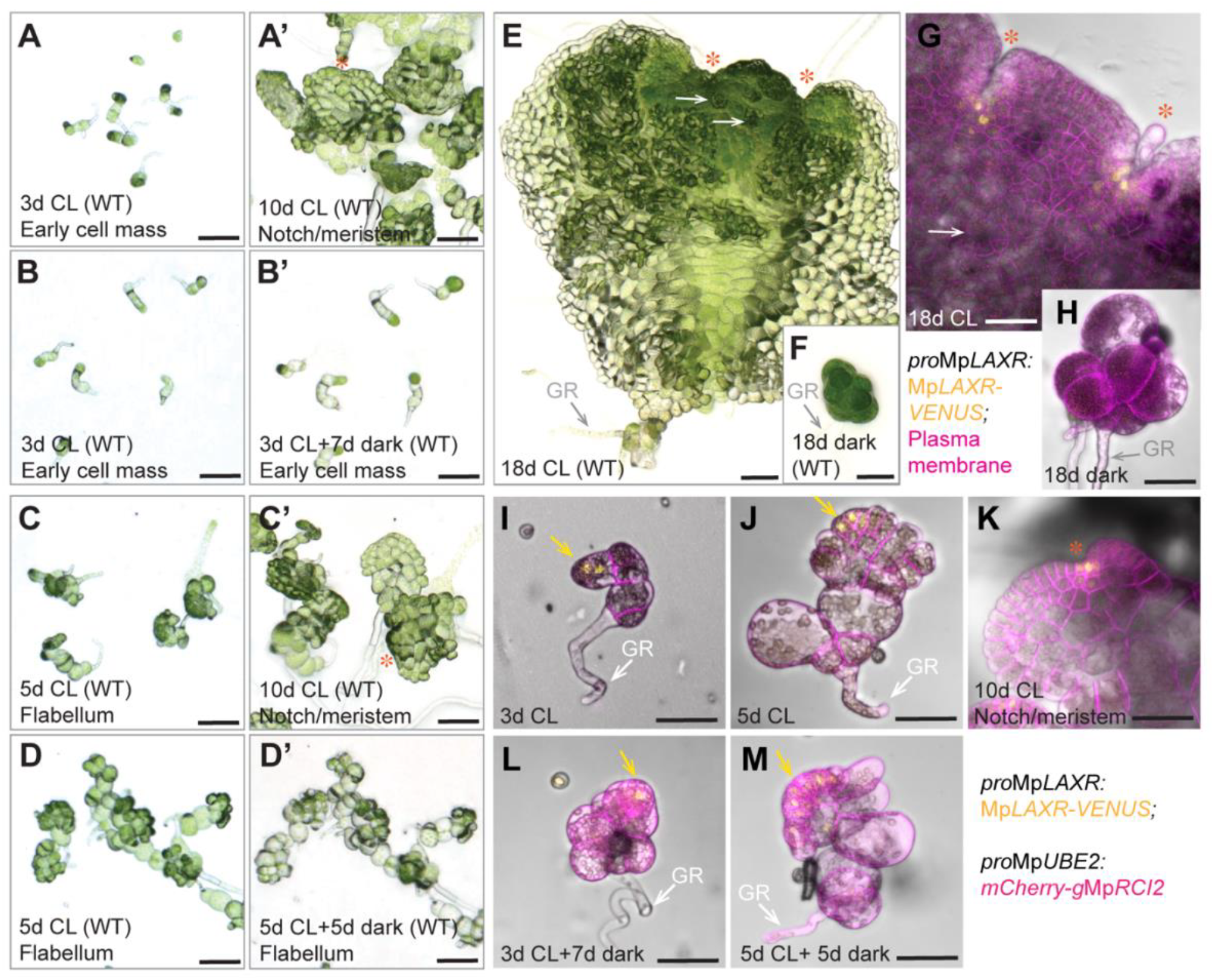
Prothallus and meristem development are light-dependent. **A-D**) Tak-1xTak-2 (wild type/WT) sporelings were grown in continuous white light (CL) for 3 (A-B) or 5 (C-D) days (abbreviated with d) and transferred to dark conditions for 7 (B’) or 5 (D’) days, respectively. Sporeling development was stopped upon transition to darkness, while sporelings grown for 10 days in CL (A’ and C’) developed the first notches (orange asterisks). Experiment was repeated 3 times with n >200. Scale bars 100 µm. **E-F**) Tak-1xTak-2 (wild type/WT) sporelings grown for 18 days in continuous white light with bifurcated meristems (orange asterisks) and differentiated thallus-like tissues, such as air pores (white arrows). Sporelings grown for 18 days in dark did not develop past the early cell mass (F). n >100, 2 repetitions. Scale bars 100 µm. **G-K**) Confocal images of sporelings expressing the transgenes *pro*Mp*LAXR:*Mp*LAXR-VENUS* (yellow signal); *pro*Mp*UBE2:mCherry-*Mp*RCI2* (plasma membrane signal in magenta) grown in continuous white light (CL) for 18 days (n = 7) showed a bifurcated meristem and MpLAXR-VENUS signal restricted to the meristems (orange asterisks) and developed the first dorsal air chambers (white arrow), while sporelings grown for 18 days in darkness (n = 53) did not develop past the early cell mass stage and were devoid of MpLAXR-VENUS signals. The germ rhizoid is marked by a grey arrow. Scale bars 50 µm. **I-M)** *pro*Mp*LAXR:*Mp*LAXR-VENUS* (yellow signal); *pro*Mp*UBE2:mCherry-*Mp*RCI2* (plasma membrane signal in magenta) expressing sporelings were grown in continuous white light (CL) for 3 (n= 24 in I), 5 (n = 39 in J) and 10 (n= 5 in K) days with MpLAXR-VENUS (yellow) located in 1-2 nuclei at the sporeling apex. After transfer to darkness for 7 (n = 21 in J) and 5 (n = 45 in K) days, development stopped and the MpLAXR-VENUS signal was maintained. Data was acquired in 5 independent experiments. Scale bars 100 µm.

To confirm that Mp*LAXR* expression marked an apical auxin minimum during sporeling development (Figure 3), we treated the *pro*Mp*LAXR:*Mp*LAXR-VENUS; pro*Mp*UBE2:mCherry-*Mp*RCI2* translational fusion line with 5 µM NAA and imaged fluorescence. While the MpLAXR-VENUS fluorescence was detectable in cells within the notch of 8-day-old mock-grown sporelings (orange asterisks in Figure 4J), no consistent MpLAXR-VENUS fluorescence could be observed in 8-day-old sporelings germinated on 5 µM NAA (Figure 4K). Likewise, transfer of 4-day-old mock-grown sporelings to 5 µM NAA for 4 days abolished apical MpLAXR-VENUS signals (Figure 4L). These data are consistent with the conclusion that MpLAXR-VENUS marks an apical auxin minimum and that formation of this auxin minimum can be disrupted by NAA treatment. NAA-treated sporelings formed cell masses comprising multiple, small and isotropically growing cells, which could be visualised by the magenta plasma membrane signals (Figure 4J’-L’). This suggests that the early cell mass can develop in the presence of high auxin, but an apical auxin minimum is required to initiate and maintain development of the prothallus. We therefore conclude that MpLAXR marks an apical auxin minimum that is first established in the early cell mass to initiate prothallus development.

### Prothallus and meristem development are light-dependent

Since the early cell mass of the sporeling grows toward the light source^4^ we reasoned that light might define the position of the apex within the early cell mass and be involved in establishing the associated local auxin minimum. To test this hypothesis, we grew wild-type spores (derived from a cross between Tak-1 x Tak-2) in continuous light and transferred them to darkness at different stages of development. Sporelings grown in continuous light for 3 days to the early cell mass stage (Figure 5A-B) and 5 days to the plate to young flabellum stage (Figure 5C-D) were transferred to darkness for a further 7 (Figure 5B-B’) and 5 (Figure 5D-D’) days, respectively. The early cell masses that were transferred to the dark at day 3 for 7 days did not initiate a prothallus (Figure 5B’). The young prothalli that were transferred to the dark at day 5 for 5 days did not expand to form a flat flabellum with meristem (Figure 5D’). By contrast, control sporelings grown in continuous light under otherwise identical conditions each developed an expanded flabellum with a notch by day 10 (orange asterisks in Figure 5A’ and Figure 5C’). By day 18, the meristems of light-grown sporelings had bifurcated and the first differentiated thallus tissues could be observed (Figure 5E). However, germinating spores and keeping them for 18 days in darkness arrested their development at the early cell mass stage (Figure 5F). We conclude that sporelings require light to transition from the early cell mass to the initiation of the prothallus and from the expanded prothallus (flabellum) to the formation of a meristem/notch (Figure 1A).

To investigate if light was also required for the formation of the apical auxin minimum marked by MpLAXR (Figure 3), we germinated *pro*Mp*LAXR:*Mp*LAXR-VENUS;pro*Mp*UBE2:mCherry-*Mp*RCI2* expressing plants in continuous light or darkness (Fiure 5G-H). After 18 days, MpLAXR-VENUS marked the notch region of the bifurcated meristem (orange asterisks in Figure 5G), while no MpLAXR-VENUS signal was observed in plants grown in darkness (Figure 5H). These data demonstrated that establishing the auxin minimum required light. To test if light was also required to maintain the apical auxin minimum and subsequently Mp*LAXR-VENUS* expression throughout development (Figure 3), we transferred MpLAXR-VENUS sporelings grown in continuous white light to darkness at the early cell mass stage (Figure 5I) and the young flabellum stage (Figure 5J). MpLAXR-VENUS was restricted to the apical stem cell in the 10-day-old light-grown control sporelings (Figure 5K). After transfer to darkness, MpLAXR-VENUS was still detectable at the apex (Figure 5L) and throughout the young flabellum (Figure 5M) even though growth and development had ceased. These data indicate that light promotes the initiation of an apical auxin mimimum but is not required for its maintenance in the short term.

### Isolated differentiated cells form early cell masses with apical auxin minima during meristem (re)generation

Removing individual *Marchantia* cells from their tissue context by surgically isolating or protoplasting them allows regeneration of whole plants without the addition of external hormones or growth factors^10,12,13^. We asked if thallus cells that regained the ability to form new meristems after surgical separation from surrounding differentiated neighbouring cells underwent a similar developmental program as sporelings. To compare the potential mechanisms that generate plants from spores with those originating from isolated thallus cells, we generated small tissue fragments from 3-day-old Tak-1 gemmalings, which were made up of young thallus that had developed from gemmae (vegetatively produced propagules)^1^. We laser ablated all but few individual cells per tissue fragment and recovered the remaining living cells within these tissue fragments on agar supplemented with standard *Marchantia* growth medium. By three days after ablation (3dpa) and recovery in white light conditions, some of these isolated cells had survived and divided to form proliferating populations of small cell clusters resembling the early cell masses of sporelings (Figure 6A-A’). Within 4-5 days a flabellum-like structure developed from each regenerant cell mass (Figure 6B-D). Approximately 7 days after ablation, the first notch formed at the apex of this flabellum-like structure (Figure 6E-E’), just as observed in the sporeling flabellum (Figure 2). To test if an auxin minimum was established during the development of meristems from individual thallus cells, we isolated single cells or tiny cell clusters from 3-day-old gemmalings expressing *pro*Mp*LAXR:*Mp*LAXR-VENUS*. Three days after ablation, these cells formed small cell masses with rhizoids that did not express MpLAXR-VENUS, like sporelings (Figure 6F). However, by day 4, MpLAXR-VENUS fluorescence was restricted to few surface cells that formed at the apical region of the proliferating cell mass of the regenerant (Figure 6G). These structures were morphologically similar to the MpLAXR-VENUS expression pattern in the plate stage of the sporeling (Figure 3D). In the young flabellum, MpLAXR-VENUS was expressed along the flabellum margin (Figure 6H) and became progressively restricted to 1-2 cells at flabellum apex (Figure 6I) and finally to 1-2 cells in the sinus of the notch (Figure 6J). This MpLAXR-VENUS expression pattern – at the flabellum apex and sinus of the notch – is the same as that observed during sporeling development (Figure 3F-H and Supplementary video 2). Together, these data suggest that similar regulatory programs that include the formation of an auxin minimum operate during the *de novo* development of meristems in sporelings derived from spores and in cell masses derived from isolated differentiated thallus cells.

**Figure 6:**
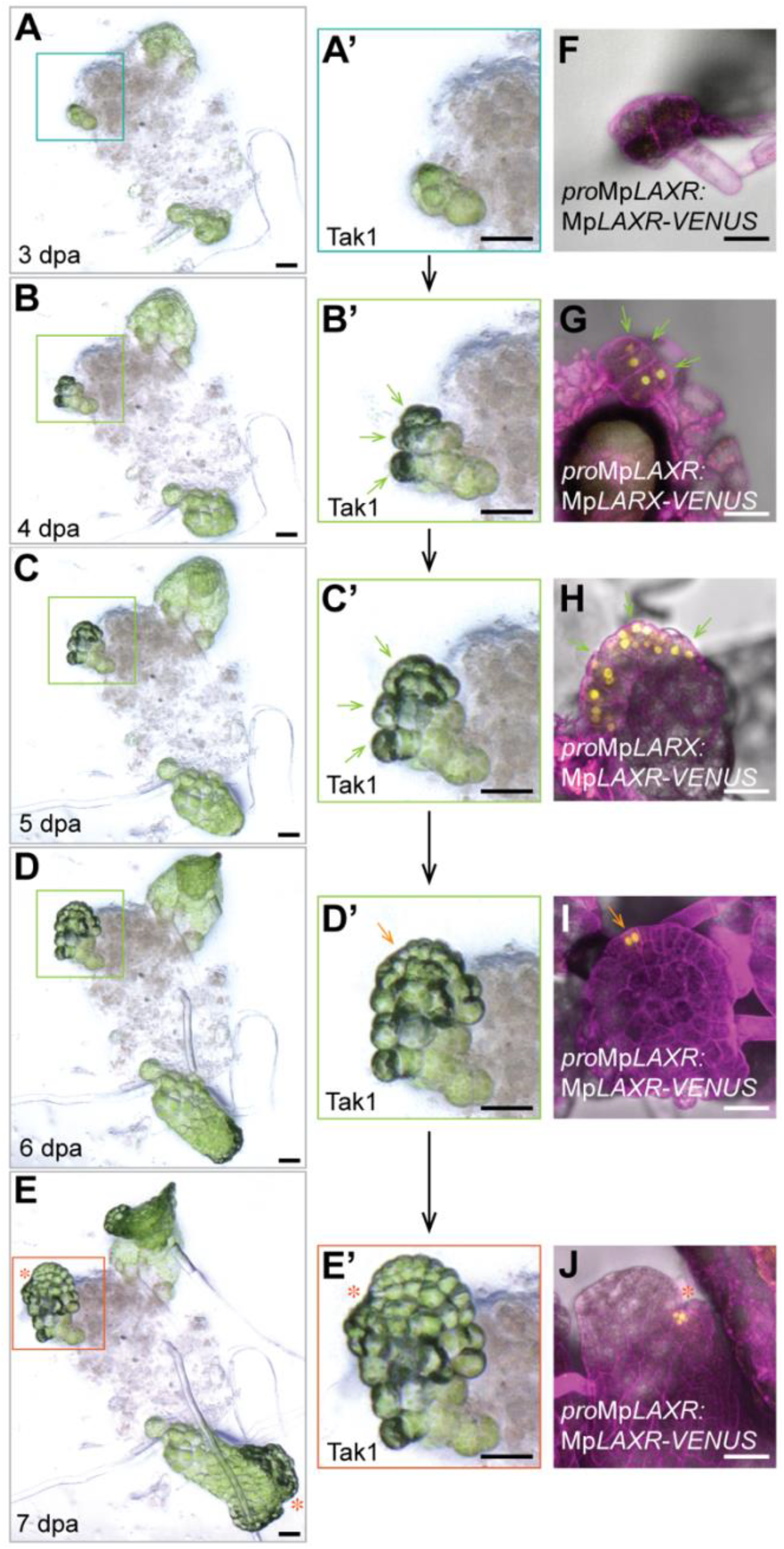
Isolated differentiated cells form early cell masses with apical auxin minima during meristem (re)generation. **A-E**) Tissue of germinated Tak-1 gemmae was chopped and ablated to obtain single living somatic cells. Bright field images were taken 3 days post ablation (dpa), where the cells had divided into early cell masses (A). The first flat prothallus structures emerged around 4 dpa (B), expanded into a round flabellum by 5-6 dpa (C-D) and formed a notch at day 7 (E). n = 100 across 3 independent experiments. Scale bars 50 µm. **F-J**) Tissue of *pro*Mp*LAXR:*Mp*LAXR-VENUS* expressing gemmae was chopped and laser ablated into single somatic cells and development of new prothalli was monitored over the course of 7 days, starting at 3 days post ablation (dpa). Tissue was stained with propidium iodide (PI, magenta cell wall signal) prior to confocal imaging. No MpLAXR-VENUS signals could be detected 3 dpa (F), but MpLAXR-VENUS signals were present throughout development of the first flat prothallus structures (G-I) and became restricted to the *de novo* formed meristem embedded in the notch (J). Scale bars 50 µm.

## DISCUSSION

Meristems contain stem cells from which multicellular plant bodies develop^19^. Although the molecular mechanisms that maintain meristem function and maintenance are well known^9,20-22^, little is known about the mechanism of meristem formation *de novo*^23^. Here we report that an external light signal establishes an internal apical auxin minimum, which promotes prothallus formation and the *de novo* development of a meristem during *Marchantia* sporeling development. We demonstrate that the same mechanism operates during the regeneration of a meristem from a single, isolated thallus cell.

Light is essential for sporeling development and initiation of the prothallus from the early cell mass. Leitgeb described that some cells of the early cell mass, which he designated “Keimschlauch” (the German word for “germ tube”), are oriented toward the light^4,24^. Furthermore, O’Hanlon showed that distinct early cell mass morphologies developed in different light intensities^5^. However, the role of light in the formation of the prothallus and the first meristem is undescribed. We discovered that light is essential for the initiation of an apical auxin minimum required for anisotropic sporeling growth and formation of the prothallus and meristem. However, light was not required to maintain the existing auxin minimum. Consistent with our observation that an auxin minimum develops at the sporeling apex is the report that PIN-FORMED (PIN) proteins, which facilitate and direct auxin transport in land plants^25^, accumulate in young *Marchantia* prothalli apices before meristem and notch formation^17^. This suggests that PIN proteins facilitate auxin efflux from the apical pole before or around the time of apical stem cell specification. This would promote the formation of an apical auxin minimum marked by Mp*LAXR* expression. Consistent with the requirement for a PIN-dependent auxin minimum in prothallus and meristem development is the observation that sporelings fail to develop beyond the early cell mass stage in Mp*pin1* mutants^17^. Combining these data, we hypothesise that light induces the formation of an apical auxin minimum to promote prothallus initiation at the apical pole of the early cell mass. This apical auxin minimum is also required later for a series of developmental transitions that occur in the prothallus leading to the initiation and subsequent development of a meristem.

The apical stem cell located in the sinus of the notch forms the centre of the *Marchantia* meristem. The timing of apical stem cell specification as well as the number of apical stem cells are still debated in contemporary literature^5,8,26,27^. Two reports suggested that the apical stem cell is derived from a row of multiple stem cells^5,26^ while another proposed that the prothallus is derived from a single stem cell^4^. Neither classical histology^4,5^ nor time lapse image analysis^8^ has established when the apical stem cell of the first *Marchantia* meristem is specified. Here we discovered that a MpLAXR-VENUS-positive cell is present at the apical pole of the early cell mass stage (Supplementary video 1) with widespread MpLAXR-VENUS signals during the early prothallus stages, including plate, disc and young flabellum. These tissues are formed by transverse, periclinal or anticlinal cell divisions^8^ and not the specific two-, three- or four-cutting face divisions ascribed to *Marchantia* apical stem cells^4,24,27^. This indicates that – while an apical site of organogenesis was marked by MpLAXR-VENUS in the early cell mass – the apical stem cell was not specified yet. In the flabellum, MpLAXR-VENUS signals became gradually restricted to a marginal flabellum cell that divided with at least three cutting faces to produce daughter cells (Supplementary video 2). These daughter cells produced two lobes on either side, restricting the MpLAXR-VENUS-expressing cell to the sinus of the developing notch (Supplementary video 2). We therefore conclude that the apical stem cell is derived from a population of MpLAXR-VENUS-expressing cells just before notch formation. This population of MpLAXR-positive cells does not include all dividing cells at the flabellum margin (Supplementary video 2). Consequently, MpLAXR helped us to distinguish the population of marginally dividing cells that eventually produce the apical stem cell from cell divisions that grow the flabellum. Imaging MpLAXR expression in timelapse therefore allowed us to accurately predict the site of apical stem cell development just before the notch forms.

The Mp*LAXR* promoter is active in regions of low auxin in *Marchantia* thalli – auxin minima^10^. Auxin is produced in the apices of vascular plants^28^, mosses^29^ and liverworts, such as in *Marchantia* meristems^9,30^. Active auxin transport into shoots and thalli generates an auxin gradient along the apical-basal axis and a local auxin minimum in the apical meristem^18,31^. In vascular plants, this auxin gradient promotes meristem activity of the primary shoot and represses outgrowth of secondary shoots, a process known as apical dominance^28^. Apical dominance is conserved in *Marchantia* and regulates meristem activity during thallus branching^11,30,32^. Our data indicates that an auxin minimum marked by MpLAXR protein, is established at the apical pole of the early cell mass. This apical auxin minimum is the site of prothallus initiation. If the single prothallus is ablated, cell divisions occur throughout the early cell mass, eventually leading to the initiation of a new prothallus. This suggests that the first prothallus represses the development of other prothalli in the sporeling, which can be seen as a form of “flabellum dominance”. This is similar to apical dominance in where dominant meristems repress meristem activity during branching. It is possible that the repressive effects of flabellum dominance over other flabella decrease with increasing sporeling size, which could explain reports of multiple flabella with meristems forming in older specimen^14^. In sporelings that were no more than 10 days old, we only ever observed a single meristem near the flabellum apex. In rare cases (2/100) where two similarly shaped flabella developed, a single meristem was positioned between the two flabella and no longer at the flabellum apex. This suggests the presence of mobile signals within flabella that determine meristem positioning. We therefore speculate that initiation of the first prothallus is controlled by an auxin-dependent inhibitory mechanism reminiscent of apical dominance. This is in line with reports about competitive sites of organogenesis that were observed after altering auxin production in gemma tissue^11^.

Spores are totipotent cells with proliferative potential while thallus or gemma cells are differentiated and do not divide. However, the terminally differentiated thallus cells reprogram upon isolation from surrounding cells and regain pluripotency to form a new meristem^10,12,13^. Nevertheless, sporelings and regenerating thallus cells that are surgically removed or protoplasted undergo comparable stages of morphogenesis as they form a meristem *de novo*^10,12-14^. These morphological stages include formation of an early cell mass, a flat flabellum and eventually a notch with meristem. Given this similar morphogenesis, we speculated if the underlying molecular programs are comparable between sporeling development and thallus regeneration^14^. We discovered that a MpLAXR-VENUS-positive cell forms toward the light-facing (apical) site of the early cell mass stage in both sporelings and regenerants from thallus cells. We thus propose that the generation of meristems during sporeling development and regeneration from single differentiated thallus cells may utilise similar molecular programs that involve the formation of an apical auxin minimum marked by MpLAXR-VENUS.

Reports on the effects of exogenous auxins on *Marchantia* development had been controversial over the last decades^11,18^. The current view is that auxin represses Mp*LAXR* promoter activity, cell divisions and meristem regeneration from intact thallus or gemma tissue^10,14,18^. In sporelings, auxin does not affect formation of the early cell mass but also suppresses development of the meristem^14^. Likewise, blocking auxin biosynthesis, signalling^16^ or transport via MpPIN1 proteins^17^ leads to growth of early cell mass-like tissues in sporelings that are either unable or delayed in forming a prothallus. These reports support our discovery that early cell masses can grow in an auxin-independent way, while the formation of an auxin minimum at the apical pole of the early cell mass is required for organogenesis.

We conclude that an auxin minimum-requiring process operates during *de novo* meristem formation from either individual spores or reprogramming single thallus cells after surgical isolation from neighbouring cells. Light is required to establish the apical auxin minimum and to promote development of the prothallus on which a *de novo* formed meristem is positioned.

## Acknowledgements

We especially thank Magdalena Mosiolek and Katharina Jandrasits (GMI) for helping with spore transformation. Special thanks also to Irmgard Fisher from the MPL for allowing us to use the LMD6500 for our experiments. We thank Vicky Spencer and Zohar Meir for critical comments to the manuscript. We also thank the BioOptics, Molecular Biology Services, Media Lab, and Lab Support of GMI/IMBA/IMP and the VBCF Plant Sciences unit for their support. This work was funded by a grant from the Austrian Academy of Sciences to the Gregor Mendel Institute and a European Research Council Advanced Grant DENOVO-P (project no. 787613) to L.D.

## Author contributions

E.-S.W. and L.D. designed the project. N.E. did the plant crossings. E.-S.W. carried out the experiments and analysed the data. E.-S.W. and L.D. wrote the manuscript.

## STAR METHODS

Detailed information about reagents and resources are listed in the Key Resource Table.

### Key resources table

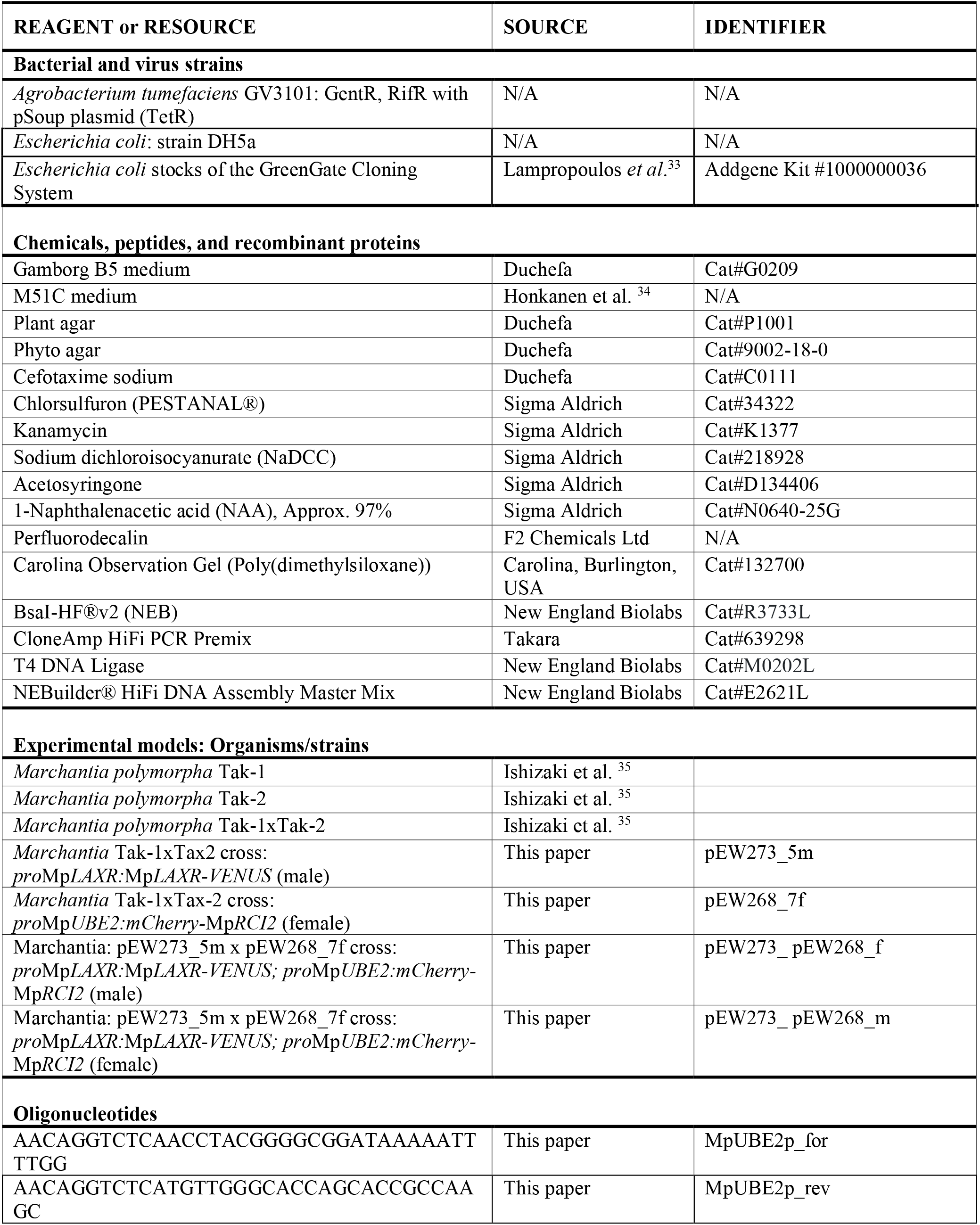
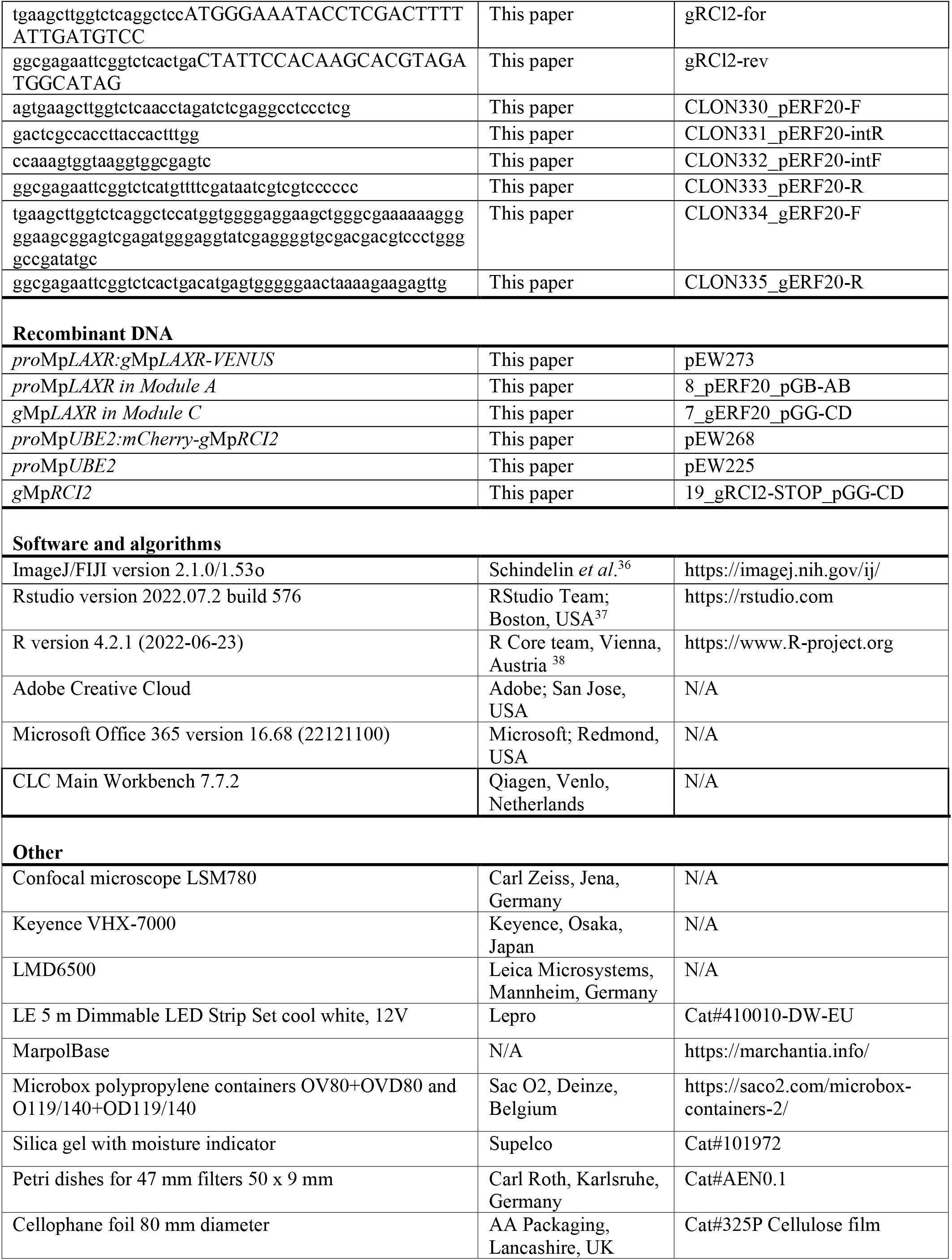

### Resource availability

#### Lead contact

Requests for resources and further information should be directed to Liam Dolan.

#### Materials availability

Newly generated lines and sources of previously reported transgenic lines are listed in the key resource table. For the transfer of transgenic material, appropriate information on import permits will be required from the receiver.

#### Data and code availability

All data required to substantiate the claims of this paper are included in main or supplemental data. This paper does not report original code.

## EXPERIMENTAL MODEL AND STUDY PARTICIPANT DETAILS

This study used *Marchantia polymorpha* wild type accessions Takaragaike-1 (Tak-1, male) and Takaragaike-2 (Tak-2, female)^35^. Plants propagated by gemma were grown on ½-strength B5 Gamborg’s agar (1.5 g/L B5 Gamborg, 0.5 g/L MES hydrate, 1 % sucrose, pH adjusted to 5.5 solidified with 1 % plant agar) at 23 °C under standard continuous white light (45 μmol m^-2^ s^-1^). To induce gametangiophores for crossing, 3 week-old gemmalings were transplanted onto an autoclaved 1:3 mixture of vermiculite and Neuhaus N3 compost and grown in SacO2 Microbox containers under 50 μmol m^-2^ s^-1^ (PPFD) white light supplemented with 45 μmol m^-2^ s^-1^ far red light irradiation in a long day regime of 16 h light, 8 h dark at 23 °C^39^.

## METHOD DETAILS

### Generation and germination of spores for sporeling analyses

Crossing of Tak-1 with Tak-2 (wild type) and *pro*Mp*LAXR:*Mp*LAXR-VENUS* (pEW273_5m) with *pro*Mp*UBE2:mCherry-*Mp*RCI2* (pEW268_7f) were performed by releasing sperm from ripe male antheridiophores in a water droplet for 5-10 minutes before transferring the liquid onto the respective female archegoniophores^39^. 3-4 weeks after crossing, sporangia formed on fertilized archegoniophores that were harvested, dried in closed microbox polypropylene containers OV80+OVD80 filled with silicagel for 5 weeks and frozen in 2 ml microcentrifuge tubes at −70°C^8^. Frozen spores were sterilised in 0.1% (w/v) sodium dichloroisocyanurate for 2-4 min, pelleted at 13000 g for 3 min and the sterilisation solution was removed and spores resuspended in 500 µl sterile water. 7 ml of ½-strength B5 Gamborg’s phyto agar (1.5 g/L B5 Gamborg, 0.5 g/L MES hydrate, 1 % sucrose, pH adjusted to 5.5 solidified with 1 % phyto agar) was solidified in small (50 mm diameter, 9 mm depth) petri dishes with safe-lock lid (Carl Roth Karlsruhe, Germany) and topped with sterilised and wetted 30 mm x 30 mm cellophane squares (#325P Cellulose film from AA Packaging, Lancashire, UK) to prevent rhizoids from entering the agar. Approximately 100 µl of sterilised spores were plated onto the cellophane squares and excess water was evaporated under sterile conditions before plates were closed. The plates were put on black paper to ensure sporeling growth toward a single light source of continuous white light (45 μmol m^-2^ s^-1^) at 23°C.

### Auxin treatments with NAA

A stock of 10 mM of 1-Naphthalenacetic acid (NAA) was prepared in dimethyl sulfoxide (DMSO) as solvent and stored at −20°C. To treat sporelings, the stock solution was added to fresh ½-strength B5 Gamborg’s agar (1.5 g/L B5 Gamborg, 0.5 g/L MES hydrate, 1% sucrose, pH adjusted to 5.5 solidified with 1% phyto agar) to a final concentration of 5 µM, mixed, poured into petri dishes and solidified before topped with sterile cellophane. For mock treatments, an equal amount of the DMSO solvent was added to the Gamborg’s agar.

### Generation of plasmids for plant transformation

Translational fusion constructs *pro*Mp*LAXR:g*Mp*LAXR-VENUS* (conferring chlorsulfuron plant resistance) and *pro*Mp*UBE2:mCherry-g*Mp*RCI2* (conferring kanamycin plant resistance) were generated using the Green Gate method^33^. The plant resistance cassette for kanamycin (pGGF007) and the mCherry tag (pGGB001) were taken from the Green Gate Cloning System kit available on Addgene (#Kit # 1000000036)^33^. The linker-VENUS module^40^ and chlorsulfuron plant resistance cassette^8^ were published previously. The sequence for *pro*Mp*UBE2* was obtained from the OpenPlant toolkit^41^ and PCR-amplified with primers MpUBE2p_for/MpUBE2p_rev (Key resource table) and the CloneAmp HiFi PCR Premix (Takara), before digested by the restriction enzyme BsaI-HF®v2 (NEB) and cloned into Green Gate Module pGGA000 with aid of the T4 DNA ligase (NEB). *g*Mp*LAXR* (Mp5g06970.1; 1083 bp genomic sequence without stop codon) and *g*Mp*RCI2* (Mp3g18270.1; 927 bp genomic sequence with stop codon) gene sequences were obtained from MarpolBase (marchantia.info) and amplified from Tak-1 genomic DNA by CloneAmp HiFi PCR Premix (Takara) with Green Gate-specific primers gRCI2-for/gRCI2-rev and CLON334_gERF20-F/CLON335_gERF20-R, respectively, as listed in the Key resource table. The amplified gene sequences were subsequently cloned via BsaI-HF®v2 (NEB) restriction sites into pGGC000. To clone *pro*Mp*LAXR*, 5000 bp upstream of the translation start site (ATG) were dedicated as promoter region and PCR-amplified in two pieces with primer pairs CLON330_pERF20-F/CLON331_pERF20-intR and CLON332_pERF20-intF/CLON333_pERF20-R using the CloneAmp HiFi PCR Premix (Takara) and assembled into the linearised pGGA000 backbone according to the NEBuilder® HiFi DNA Assembly Master Mix (NEB) manual. Final constructs were assembled using entry modules and the pGreen-IIS based vector pGGZ003 as backbone according to the Green Gate method^33^. Correct plasmid assembly was confirmed by full plasmid sequencing at the Microsynth AG.

### Confocal and time lapse imaging of sporelings

Confocal images were acquired on an LSM 780 system (Carl Zeiss) with an upright 20x/0.8 plan-apochromat air objective. Fluorescent fusion proteins VENUS and mCherry were excited by an argon laser of 514 nm and a DPSS diode of 561 nm, respectively. Emission spectra were collected between 520-540 nm (VENUS) and 600-650 nm (mCherry) in the sequential scanning mode. A self-made time lapse chamber was built based on instructions by Kirchhelle and Moore that used a glass casket made out of objective slides that was filled with perfluorodecalin and sealed with air-permeable gas-permeant poly(dimethylsiloxane) gel^42^. An in-detail description of time lapse chamber assembly that was optimised for the use of sporelings was previously published^8^. Time lapses were acquired using the VENUS and/or mCherry settings with 1 hour intervals in-between frames. The frames were assembled and exported as video file using ImageJ/FIJI.

### Generation and origin of transgenic lines

Transgenes were introduced into wild type sporelings generated from a cross between Tak-1 and Tak-2 wild-type accessions of *Marchantia polymorpha*. The suspension of one sterilized sporangium was cultured in sterile 125 ml Erlenmeyer flasks with 25 ml M51C medium^35^ for 7 days at 23 °C under continuous white light (45 μmol m^-2^ s^-1^) with constant agitation (130 rpm). Electrocompetent *Agrobacteria* (see key resource table) were transformed with the desired plasmid and colonies were picked to grow a dense overnight culture in 5 ml LB-medium with respective antibiotics. The *Agrobacteria* culture was pelleted and the pellet resuspended in 10 ml M51C medium ^34^ containing 100 mM acetosyringone and incubated for 6 h at 28°C. The 7-day old sporeling culture was mixed with 1 ml of the *Agrobacterium*-induced M51C culture and co-cultivated for 2 days with a final concentration of 100 mM acetosyringone. The sporeling culture was washed with 250 ml sterile water using a 40 mm nylon cell strainer and plated on ½-strength B5 Gamborg’s agar (1.5 g/L B5 Gamborg, 0.5 g/L MES hydrate, 1% sucrose, pH adjusted to 5.5 solidified with 1% plant agar) with 10 mg/ml cefotaxime and 0.5 µM chlorsulfuron (pEW273) or 50 µg/ml kanamycin (pEW268) for selection. Crosses between *pro*Mp*LAXR:*Mp*LAXR-VENUS* and *pro*Mp*UBE2:mCherry-g*Mp*RCI2* were used for confocal imaging and time lapses.

### Sporeling phenotyping and cell ablations

Wild-type and mutant plants were imaged with the Keyence VHX7000 (Keyence, Osaka, Japan) digital microscope equipped with the VH-Z00R/Z00T and VH-ZST lenses and the VHX-7020 camera. For cell ablations, small petri dishes with sporelings were opened under a Leica Laser Microdissection System LMD6500 (Leica, Mannheim, Germany) and cells were ablated with the 355 nm DPSS laser set to 60 power, 10 aperture, 17 speed using a 20x/0.4 PL-Fluotar correction collar objective. Images of sporelings before and after ablations were taken at the Keyence VHX7000.

### Regeneration from single somatic gemma cells

For regeneration studies from single somatic cells, gemma were germinated on 0.5x Gamborg agar plates on sterilized cellophane and chopped in 300 µl sterile water with a razor blade into small pieces. The tissue pieces were spread onto a fresh 0.5x Gamborg agar plate and all cells except for single somatic cells at the outmost edge were ablated under a Leica Laser Microdissection System LMD6500 (60 power of 355nm DPSS laser, 10 aperture, 17 speed under the 20x/0.4 PL-Fluotar correction collar objective). Tissue pieces were recovered in a light chamber with standard continuous white light for 3 days, in which many cells died. The surviving isolated somatic cells were tracked from 3 days post ablation onwards using the Keyence VHX7000.

## QUANTIFICATION AND STATISTICAL ANALYSIS

Microscopy images were analysed in ImageJ/FIJI version 2.1.0/1.53o^36^ and measurements of apical-basal distance and circularity and the angles of notch positioning the oval, straight and angle measurement tools were used. Measurements were statistically analyzed in R^38^ and Rstudio^37^ version 2022.07.2+576 and plots were created with the ggplot2^43^ package. Other packages used: tidyverse, ggbeeswarm, scales, ggpubr, knitr, ggsignif, FSA, ggstatsplot, rcompanion and multi-compView (for detailed information see www.r-project.org). Datasets were first tested on normal distribution by a Shapiro-Wilk Test and homogeneity of variances by a Levene-Test (α=0.05). A one-sample t-test (p-value <0.05) was performed to test how much meristem positioning deviated from 90° and data was presented in a scatter and violin plot. To determine flabellum positioning within the sporeling, an ellipse was fitted to encompass all green sporeling cells using ImageJ/Fiji. The lengths of major axes and minor axes were determined along the apical-basal body axis, which is the distance between primary rhizoid base and flabellum apex. The ratio of minor to major axis of the fitted ellipse was compared to the ratio of apical-basal to the major axis using a paired Wilcoxon test (p-value <0.05) and plotted as scatter, box and violin plots. The precision measures of all box plots are medians flanked by lower and upper hinges corresponding to the 25^th^ and 75^th^ percentile, respectively. Box plot whiskers extend from each hinge to the largest and smallest value measured with maximal 1.5-times the inter-quartile range. The type of statistical test used, p-values and sample sizes (n) are indicated in the figure legends. Precision measures of all other types of plots are described in the respective figure legends. Each statistical analysis presented displays one of at least 2-9 independent replicates.

### Video S1. MpLAXR-VENUS marks an auxin minimum that is specific to the most apical sporeling cell

Confocal time lapse imaging depicts a *pro*Mp*LAXR:*Mp*LAXR-VENUS* (yellow signal) expressing sporeling imaged with the transmitted light channel to depict cell outlines. 23 frames with 1 hour intervals were taken between day 4 and day 5. Scale bar 50 µm.

### Video S2. MpLAXR-VENUS marks the future apical stem cell prior to notch formation

Confocal time lapse imaging depicts a *pro*Mp*LAXR:*Mp*LAXR-VENUS* (yellow signal); *pro*Mp*UBE2:mCherry-*Mp*RCI2* (plasma membrane signal in magenta) expressing sporeling imaged for 55 hours (between day 7 and day 10) with frame intervals of 1 hour. Scale bar 50 µm.

